# Measuring narrative engagement: The heart tells the story

**DOI:** 10.1101/351148

**Authors:** Daniel C. Richardson, Nicole K. Griffin, Lara Zaki, Auburn Stephenson, Jiachen Yan, Thomas Curry, Richard Noble, John Hogan, Jeremy I. Skipper, Joseph T. Devlin

## Abstract

Stories play a fundamental role in human culture. They provide a mechanism for sharing cultural identity, imparting knowledge, revealing beliefs, reinforcing social bonds and providing entertainment that is central to all human societies. Here we investigated the extent to which the delivery medium of a story (audio or visual) affected conscious and subconscious engagement with the narrative. Although participants self-reported greater involvement for watching video relative to listening to auditory scenes, stronger physiological responses were recorded for auditory stories. Sensors placed at their wrists showed higher and more variable heart rates, greater electrodermal activity, and even higher body temperatures. We interpret these findings as physiological evidence that the stories were more cognitively and emotionally engaging when presented in an auditory format. This may be because listening to a story, rather than watching a video, is a more active process of co-creation, and that this imaginative process in the listener’s mind is detectable on the skin at their wrist.

---

Stories help us make sense of the world. Narratives provide links to traditions, legends, archetypes, myths, and symbols and help connect us to others by forming and stabilizing social bonds, by reinforcing and enhancing the group’s memory, and by providing shared entertainment. Our oldest narratives date back many thousands of years and pre-date the advent of writing. For the majority of human history, stories were synonymous with the oral tradition; audiences listened to a story teller imparting a tale. In modern cultures, stories are just as important but now are delivered in a variety of mediums including written books (both physical and digital), videos (TV and films), and of course, as auditory narratives. Here we investigated the extent to which the delivery medium (audio or visual) affected one’s conscious and sub-conscious engagement with the narrative.

A good story takes the listener on a journey, evoking cognitive and emotional responses such that the listener experiences the story through a process of mental simulation of the people, events, actions, places and emotions from the narrative, as if these were being experienced directly. Indeed, there is evidence that narratives recreate a similar pattern of brain activity in the listener that was produced by the storyteller. Silbert and colleagues (2014) used functional magnetic resonance imaging (fMRI) to scan the brain of a volunteer speaking a 15-minute personal story. Another set of volunteers then listened to this story while having their brains scanned. The authors identified the set of brain regions engaged by these tasks and found widespread coupling between activity in the speaker’s brain and that in the listeners’ brains. In other words, the act of listening to the narrative recreated the same basic pattern of brain activity as telling the story, suggesting that listening to the story is qualitatively and quantitatively similar to experiencing the speaker’s memory of the events. Moreover, activation was not limited to regions of the brain classically related to language, but also involved emotional, sensory and motor systems consistent with the notion that at some level, the listener actually experiences the story.

Historically, story-telling relied primarily on spoken language, and then more recently on written language, but in the modern era video has emerged as a major narrative tool as well. The main difference between these channels is the amount of information they provide. Spoken words come in a single modality, namely audition, and have a very abstract relation to the content of the narrative. Consider a story that contains the sentence: “The was house ablaze.” A listener will correctly interpret this to mean that the house was on fire and possibly imagine what it might be like but the actual physical stimulus – in this case, changes in acoustic energy over time – is unrelated to the content being conveyed except through the interpretation of language. Video, on the other hand, is more closely related to the content. Seeing a video of a burning house, hearing the sounds of the fire – these are physical stimuli that directly convey the information without interpretation and without language. Consequently, there is a very different level of engagement between these channels. Oral and written stories require a more active engagement as the listener/reader reconstructs a personalized interpretation of the narrative. In contrast, watching video is a more passive process due to the fact that there is less scope for personal interpretation.

The question we asked here is whether a difference in the delivery channel would manifest as differences in engagement with the narrative, measured by a combination of explicit self-report and implicit physiological measurement. Participants listened to auditory and watched video scenes chosen from well-known novels with audio and video adaptations. They were asked to rate their experience of scene using a narrative engagement instrument. In addition, biometric sensors were used to measure heart rate, electrodermal activity and body temperature. Changes in heart rate have been linked to increased information processing demands and/or greater mental effort (Andreassi, 2007; Potter & Bolls, 2012; Sukalla, Bilandzic, Bolls, & Buselle, 2016) while both electrodermal activity and body temperature have been associated with emotional arousal (Critchley, 2002). All of these physiological signals are also affected by physical activity, and so we used accelerometers on the sensors to track and account for body motion.

## Method

### Participants

109 participants were recruited from UCL’s subject pool and paid £10 for participation. 7 participants were excluded due to equipment failure or participant drop out. Of the 102 (41M, 61F) who completed the experiment, their ages ranged from 18 to 55 with an average age of 29 years old (SD = 10.5). The sensors we used sometimes failed to record complete data, due to problems with the sensor, incorrect placement, the participant adjusting the wristband, and so on. The failure rate differed for different sensors as they were more or less susceptible to interruption and artifacts. After identifying and cleaning problematic recordings, we were left with 95 participants with complete heart rate and acceleration data, 76 with temperature data, and 62 with complete electrodermal activity data.

### Stimuli and Materials

We chose eight works of fiction spanning four different genres. From classic literature we selected *Pride and Prejudice* (Austen, 1797) and *Great Expectations* (Dickens, 1860); from action novels we chose *The Girl on the Train* (Hawkins, 2015) and *The DaVinci Code* (Brown, 2003); from crime fiction we chose *Hound of the Baskervilles* (Conan Doyle, 1887) and *The Silence of the Lambs* (Harris, 1988) and from Science Fiction/Fantasy we chose *Alien* (Golden, 2014) and *A Song of Ice and Fire* (Game of Thrones) (Martin, 1991). Each of these was selected because they were well-known examples of their genres and because audio and video adaptations of the novel were available. For each, we selected an emotionally charged scene where the audio and video versions were as similar as possible. For example, from the *Game of Thrones* book (Martin, 1991) we chose the passage in which Arya witnesses her father’s beheading, and the same scene in the HBO adaptation (Benioff & Weiss, 2015). Though they covered the same events, due to their nature, the audiobooks tended to be slightly longer (M=398 secs, SD=210) than the video versions (M=295 secs, SD=117).

To measure how much participants were engaged in the stories, we adapted the narrative engagement scale developed by Busselle and Bilandzic (2009), which has been validated and shown to be related to physiological measures (Sukalla et al., 2016). The scale is divided into four subscales with three questions relating to each: emotional engagement (e.g. “I understand why the main character thought and behaved as they did in the story”), narrative understanding (e.g. “I had hard time recognizing the thread of the story”), attentional focus (e.g. “I had a hard time keeping my mind on the story”), and narrative presence (e.g. “At times, the story was closer to me than the real world”).

### Procedure

Participants were informed about the experiment and gave their consent to take part. They were fitted with an Empatica E4 wrist sensor, which captured their heart rate (HR), electrodermal activity (EDA), acceleration in 3 dimensions, and wrist temperature. They were led into a sound attenuated cubicle. The experiment was run on a PC, using the Gorilla online testing platform (https://gorilla.sc), and participants wore headphones throughout. They first completed a short demographic questionnaire and a survey asking about their consumption of movies, books and audiobooks.

In the main experiment, participants were presented with 8 stories, in a block of four audio books and four videos. Across participants we counterbalanced the order of these blocks, and whether a particular story was presented as a video or audio book.

In each trial, the participant first read a short synopsis of the plot and characters in the story so far, to give a context for the excerpt. They then watched the video onscreen or listened to the story while looking at a black screen. After presentation, participants reported whether or not they had experienced that excerpt before or not, and dragged a slider to indicate how familiar the characters were, and how familiar the scene was. Then they rated the 12 statements of the narrative engagement questionnaire, using a 7 point Likert scale that ranged from “strongly disagree” to “strongly agree.” The experiment took approximately an hour to administer. On completion the participants were debriefed, thanked and paid for their time.

### Data Processing

Physiological data were aligned to stimulus and condition information and trimmed to trial durations using the Universal Time Coordinates that were recorded by the Empatica sensors and the Gorilla system. EDA measurements are typically susceptible to movement artifacts, and so we used the EDA Explorer algorithm (Taylor et al., 2015) to clean the EDA data using the acceleration data. The 3-dimensional acceleration vectors were then simplified into a single acceleration value that expressed movement magnitude in any direction.

All physiological measures were normalised for each participant. This removed any baseline differences between individuals (e.g. their resting heart rates) and allowed us to focus on differences between story modalities within each participants’ data. The data are available from the Open Science Foundation at https://osf.io/u452g/.

## Results

Participants reported that the videos were more engaging than the audiobooks by about 15% on average across our measures. Conversely, participants’ physiological measures showed higher engagement for audiobooks rather than videos. In terms of raw measures, their average heart rate was higher when they were listening to audiobooks by about two beats a minute; they had a greater range of heart rate by about 4 beats per minute; they were roughly a third of a degree warmer in their body temperature (0.34°C), and their skin conductance (EDA) was higher by 0.02 microsiemens.

Figure 1 shows an example of the time-course of our physiological measures for the *Game of Thrones* story in two modalities. Since the audiobook and video had different durations, we have plotted our measures as a function of the proportion of the story time. The differences shown for this item were echoed across all stories (see supplementary materials).

**Figure 1.**
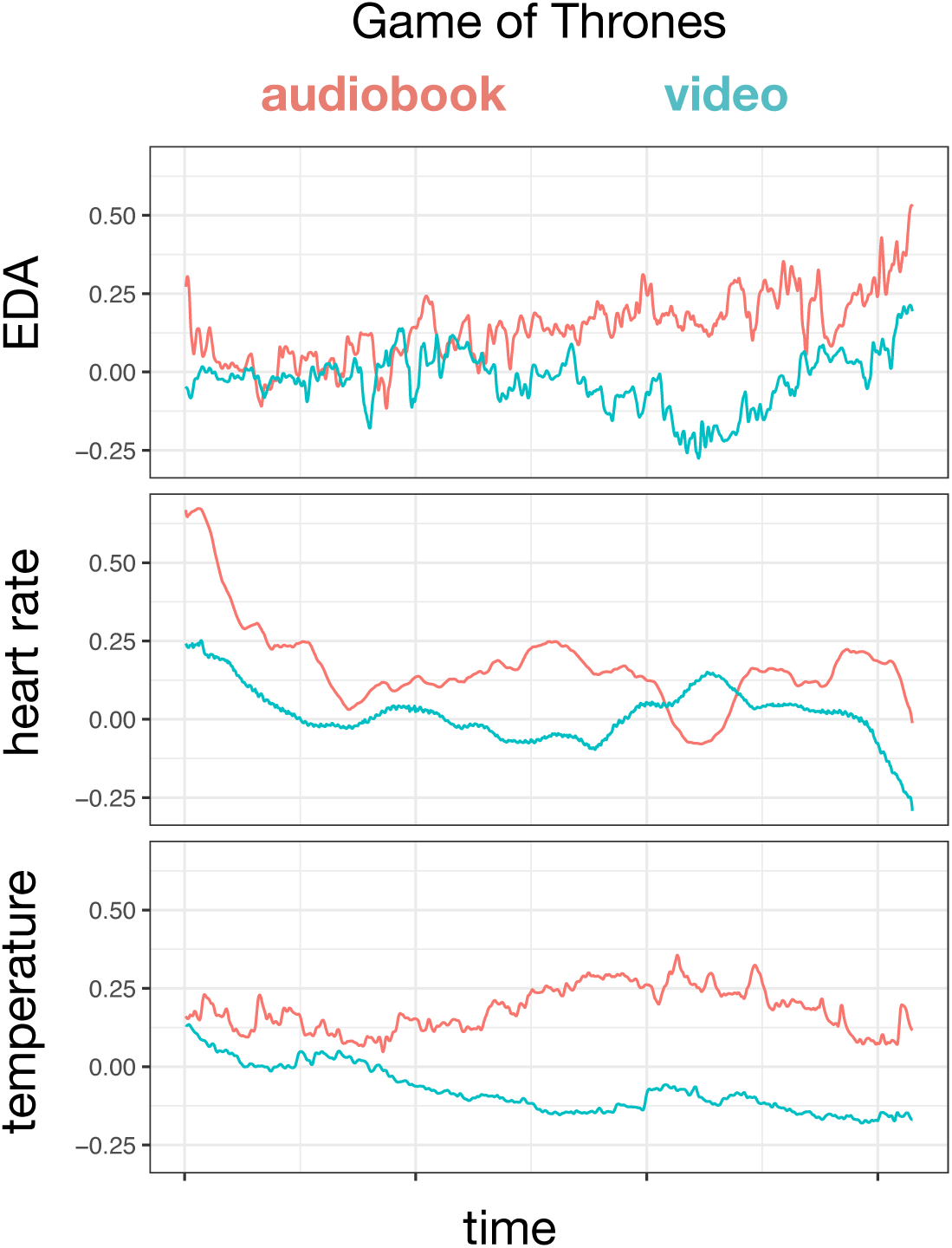
Time-course of physiological measures during the Game of Thrones story in two modalities.

Figure 2 presents the means and distributions for the participants’ engagement ratings and normalized physiological measures, contrasting audio and video modalities. Beneath the observed data are probability distributions for the estimated differences between modalities. These estimates were derived from the posterior distributions given by Bayesian mixed models of our data (Sorensen, Hohenstein, & Vasishth, 2016).

**Figure 2.**
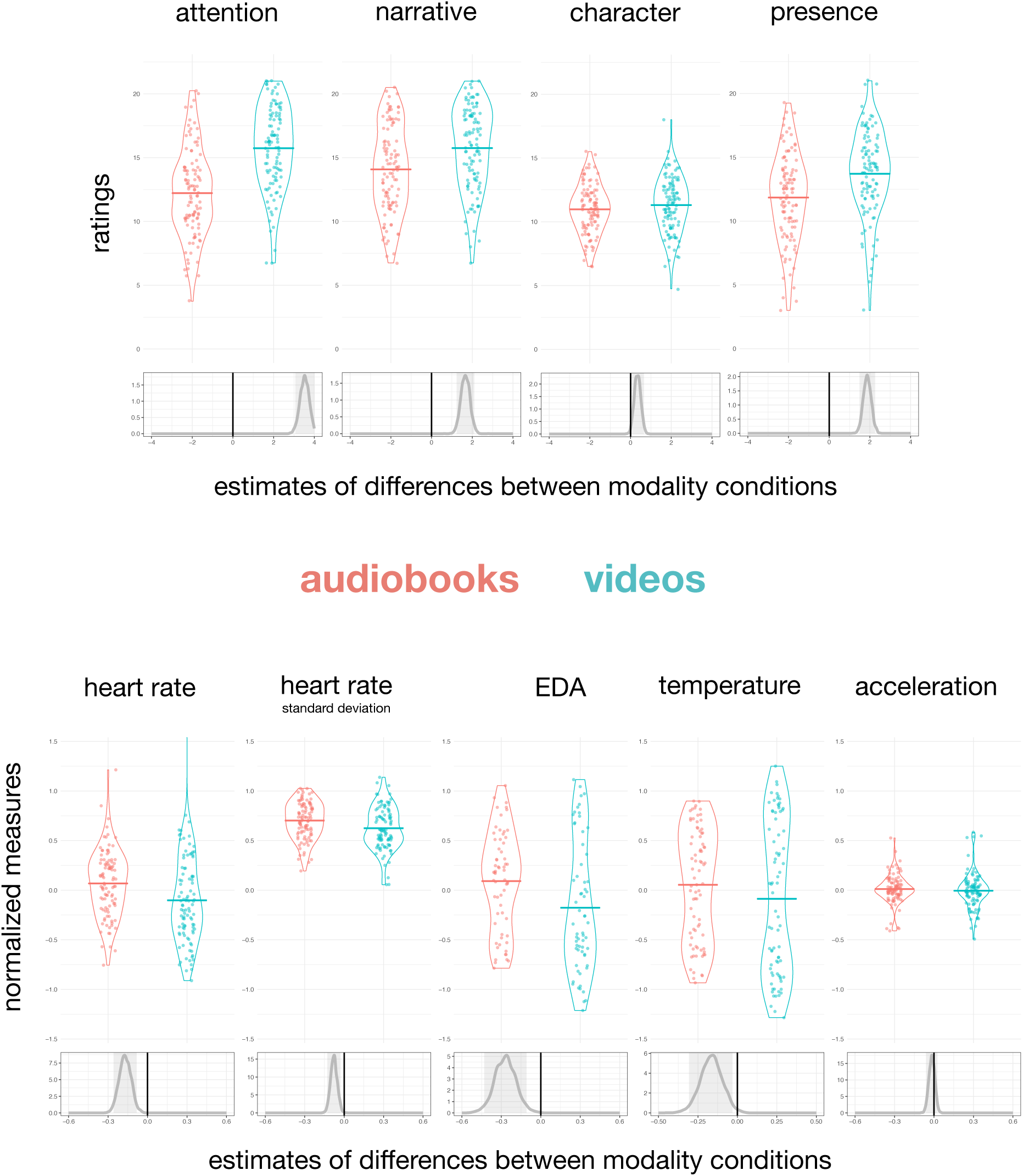
Participant engagement ratings and physiological measures, split between story modalities. Beneath observed data are probability distributions for the estimated difference between modalities, derived from Bayesian mixed models. Grey shaded areas represent 95% credibility intervals. When this region does not include the black line of zero difference, there is strong evidence supporting a difference between conditions.

The Bayesian approach allowed us to directly quantify the effects of modality on behavioural and physiological measures and the strength of evidence in support of any differences, avoiding some of the problems associated with null hypothesis testing (Kruschke, 2010; Wagenmakers, Wetzels, Borsboom, & van der Maas, 2011). In the supplementary information, we also report more traditional ANOVAs analyses, which produced the same pattern of conclusions.

For each of the dependent variables reported below, we used mixed models with fixed effects for story modality, which was varied within participants, and block order and stimuli order, which were varied between participants as counterbalancing measures. We used random effects for participants, story, and trial number, to model changes in measures during the course of the experimental hour. We used R (version 3.4.3) and the *rstanarm* package (Stan-Development-Team, 2016). In our models, we employed weakly informative priors that were scaled following the standard rstanarm procedure. From 4000 simulations, we generated estimates of the posterior distributions of the model parameter coefficients, which quantify the strength of the evidence that each experimental condition influenced behaviour. Full details of our models, priors, and all parameter estimates are given in the SI. Below we report the strength of evidence in favour of differences between audio and video modalities.

### Reported Engagement

There was strong evidence that participants rated their engagement higher for videos rather than audiobooks. Summing the 3 questions for each subscale gave a value between 3 to 21. For their attention to the story, the median of the posterior distribution for the effect of the video condition was 3.5 points higher, with a 99.99% chance that it was between 2.63 and 4.44. For their engagement with the narrative, the median was 1.67 points higher, with a 99.99% chance that it was between 0.94 and 2.42. For their ratings of presence, the median was 1.88 points higher with a 99.99% chance of being between 1.14 and 2.65. For rated engagement with characters, there was a smaller difference of 0.34 points in favour of the videos, with a 98.15% that was between 0 and 1.03. In other words, there was strong evidence that participants self-reported greater attention, engagement with the narrative and presence in the story for videos relative to audio stories.

### Physiological measures

The physiological evidence consistently demonstrated stronger responses for the audio relative to the video condition. The median estimate of normalized mean heart rates was 0.17 higher for audiobooks, with a 99.99% chance of being between 0.002 and 0.34. The median standard deviation of heart rates was also greater by 0.07, with a 99.95% chance of being between 0 and 0.17. The median of the estimated EDA readings was 0.27 higher for audio books, with a 99.99% chance of being between 0.025 and 0.55, and participants’ wrist temperature had a median estimated increase of 0.17, with a 99.20% chance of being between 0 and 0.5.

One possible explanation for the higher physiological responses may be that participants were more physically active when listening to audiobooks – that is, they could have been fidgeting more, consistent with their self-reported lower engagement. To assess this, we examined the acceleration data from the Empatica sensors. There was no strong evidence that participants moved more during audiobooks, with the median estimated difference at 0.017, and only 76.20% chance that the coefficient was between 0 and .11. The lack of a difference suggests that the differences in heart rate, EDA and temperature were not due to physical movement, but instead due to the emotional and cognitive differences in listening to audiobooks.

A second potential confound between the modalities was the fact that on average, audio scenes were longer than the equivalent video clips by approximately 100 seconds. To avoid cumulative differences in effort over time, we compare mean scores. Even so, if there were an upward linear trend with time, this could potentially inflate the difference with the longer clips showing larger effects. The data from all eight stories are shown in Supplemental Figure 1. To investigate whether the longer clips affected the results, we trimmed all the physiological data to the length of the shortest modality (usually the video clip) and re-analysed the results. The point that the stories were trimmed is shown in Supplemental Figure 1. The median estimate of normalized mean heart rates was 0.15 higher for audiobooks, with a 99.98% chance of being between 0 and 0.32. The median standard deviation of heart rates was also greater by 0.04, with a 94.8% chance of being between 0 and 0.15. The median of the estimated EDA readings was 0.27 higher for audio books, with a 99.92% chance of being between 0 and 0.55, and participants’ wrist temperature had a median estimated increase of 0.19, with a 99.52% chance of being between 0 and 0.47. In other words, the pattern of findings remained the same when the clips were truncated to the same duration, even though the peak emotional point tended to be at the very end of the scene and therefore was removed from these analyses.

## Discussion

Participants reported higher levels of engagement while watching video scenes compared to listening to audio scenes. They attended more, showed greater narrative understanding and reported greater narrative presence when watching video clips, suggesting that they not only found video narratives easier to comprehend, but also immersed themselves more fully in the world created by the video narratives. In other words, people found the videos more engaging according to their self-report. Interestingly, their implicit physiological measures told a different story. On average, heart rates were higher and more variable, ectodermal activity was greater and temperatures were raised when listening to audio narratives than when watching video narratives. These findings suggest that listening to audio stories engaged greater cognitive and emotional processing than watching videos.

In its simplest form, increased heart rate is an indicator of increased effort, consistent with the hypothesis that listening to a story is a more active process than viewing the same story. In essence, the listener mentally simulates the narrative whereas the viewer passively processes the visualization provided by the video’s director. A recent study illustrated this by measuring brain activity while volunteers listened to stories that were either visually vivid, action-based, or emotionally charged (Chow et al., 2014). All three story-types activated the temporal lobes and Broca’s area, as expected, but the interesting findings pertained to the differences between the stories. Specifically, visually vivid stories activated the occipito-parietal junction and the pre-cuneus, two regions associated with visuo-spatial processing. Action-based stories, in contrast, activated regions of premotor cortex while emotionally laden stories activated parts of the limbic system typically linked to affective responses, demonstrating that listening to stories engaged not only core “language regions” of the brain such as Broca’s area, but also recruited additional brain systems depending on the context. This is consistent with the notion that understanding a narrative involves mental simulation that retrieve the listener’s perceptual, motor, and affective knowledge through reactivation of the neural systems responsible for perception, action, and emotion.

Another fundamental difference between purely language based stories and video is the presence of semantic ambiguity. Semantic ambiguity refers to the fact that most words in English have more than one meaning (Rodd, Gaskell, & Marslen-Wilson, 2002) which means that listeners/readers are frequently resolving ambiguities, often without even noticing them. For example, in a sentence like “The woman made the <u>toast</u> with a new *microphone,*” the word “toast” is ambiguous – it could refer to cooked bread or a call to drink together. It is not until the word “microphone” is encountered that the meaning becomes clear. Although this appears to occur effortlessly, resolving ambiguity is a complex process involving multiple cognitive operations, supported by a set of brain regions including Broca’s area and posterior parts of the temporal lobe (Rodd, Davis, & Johnsrude, 2005; Zempleni, Renken, Hoeks, Hoogduin, & Stowe, 2007). There is, however, no ambiguity in the video equivalent where the image of a woman speaking into a microphone is clear from the outset. As a result, even if they are unaware of it, the listener is working harder to understand the story than a person viewing a video would. As a result, listening to a story or reading it is a more active process than watching the video.

Linking heart rate to specific cognitive states, however, is not straightforward. Andreassi (2007) claimed that heart rates increase when people focus more on internal information and less on the external environment, which could suggest that participants were retrieving their own memories in response to the audio narratives. Indeed, Papillo and Shapiro (1990) claim that increased heart rate demonstrates cognitive elaboration, consistent with a more active engagement with audio stories. Others, however, have argued that higher heart rates indicate reduced emotional engagement (Jola, Grosbras, & Pollick, 2011), suggesting *less* engagement with the audio stories. In the current example, the additional physiological measures (electrodermal activity and body temperature) may help to distinguish between these possibilities.

We observed significantly higher electrodermal activity (EDA) when participants listened to audio narratives compared to when they watched the same video narratives. EDA is a typically understood as a measure of emotional arousal (Critchley, 2002; Sequeira, Hot, Silvert, & Delplanque, 2009). One of the key emotional centres in the brain, the amygdala, stimulates the adrenal medulla, releasing the hormone adrenaline and enhancing autonomic nervous system activity. One consequence is the constriction of sweat glands in the dermis which increase skin conductance. In other words, the EDA results provide an indirect demonstration that when listening to audio stories, participants experienced greater emotional arousal than when watching video stories.

The skin temperature results tell a similar story of greater engagement by audiobooks over videos. Changes in body surface temperature reflect shifts in blood flow to the surface of the skin, a process that is controlled by the autonomic nervous system. This system responds to changes in mood and social context (Ioannou, Gallese, & Merla, 2014). Since thermoregulation is biological costly, IJzerman and colleagues (2015) argue that many social animals have evolved to share body warmth between themselves by directing blood towards the skin, and then huddling or engaging in skin-to-skin contact. As a consequence, it has been found that skin temperature on the hands increases by a fraction of a degree when participants watch film clips that produced positive, happy affect (Rimm-Kaufman & Kagan, 1996) or engage in positive social interactions (Hahn, Whitehead, Albrecht, Lefevre, & Perrett, 2012), whereas Kistler et al. (1996) found deceases in finger temperature response to fear inducing stimuli.

In the current experiment, neither the narrative instrument nor the physiological measures had the resolution to clearly distinguish between cognitive and emotional engagement. Further work will be necessary to disentangle these contributions.

## Conclusion

We found a disconnect between implicit and explicit measures of engagement. Participants perceived themselves to be more concentrated and engaged while watching video narratives, but their physiological responses revealed more cognitive and emotional engagement while listening to audio narratives. Why do they feel more engaged if their bodies say otherwise?

We suggest that spoken narratives require the participant to be an actively engaged listener, whereas videos deliver rich stimulation to a passive viewer. The pictures in the listener’s mind may not be as vivid and as detailed as those onscreen, and so auditory narratives are rated explicitly as less engaging; yet the imaginative generation of those images requires greater cognitive and emotional processing, and so they are physiologically more engaging.

